# First in-human PET study of 3 novel tau radiopharmaceuticals: [^11^C]RO6924963, [^11^C]RO6931643, and [^18^F]RO6958948

**DOI:** 10.1101/296764

**Authors:** Dean F. Wong, Robert A. Comley, Hiroto Kuwabara, Paul B. Rosenberg, Susan M. Resnick, Susanne Ostrowitzki, Cristina Vozzi, Frank Boess, Esther Oh, Constantine G. Lyketsos, Michael Honer, Luca Gobbi, Gregory Klein, Noble George, Lorena Gapasin, Kelly Kitzmiller, Josh Roberts, Jeff Sevigny, Ayon Nandi, James Brasic, Chakradhar Mishra, Madhav Thambisetty, Abhay Mogekar, Anil Mathur, Marilyn Albert, Robert F. Dannals

## Abstract

**Background:** [^11^C]RO-963, [^11^C]RO-643 and [^18^F]RO-948 (previously referred as [^11^C]RO6924963, [^11^C]RO6931643, and [^18^F]RO6958948, respectively) have been reported as promising PET tracers for tau imaging based on in vitro and preclinical PET data (*1,2*). Here we describe the first human evaluation of these novel radiotracers.

**Methods:** Amyloid PET positive Alzheimer’s disease (AD) patients and young healthy subjects (YC) each received two different tau tracers. Dynamic 90 min scans were obtained after bolus injection of [^11^C]RO-963, [^11^C]RO-643 or [^18^F]RO-948. Arterial blood sampling was performed in 11 healthy controls (HC) and 11 AD. Regions were defined on MRI, and PET data were quantified by plasma reference graphical analysis (for V_T_) and target cerebellum ratio (SUVR60-90). SUVR images were also analyzed voxelwise. Five older healthy subjects (OC) each received two scans with [^18^F]RO-948 for evaluation of test-retest variability. Four AD subjects received a repeat [^18^F]RO-948 scan over about 1 year. Six additional HC (3M: 3F; 41-67y) each received one whole body dosimetry scan with [^18^F]RO-948.

**Results:** In YC, peak SUV values were observed in the temporal lobe with values of approximately 3.0 for [^11^C]RO-963, 1.5 for [^11^C]RO-643 and 3.5 for [^18^F]RO-948. Over all brain regions and subjects, the trend was that [^18^F]RO-948 had the highest peak SUV value, followed by [^11^C]RO-963, and then [^11^C]RO-643. Regional analysis of SUVR and V_T_ for [^11^C]RO-643 and [^18^F]RO-948 clearly discriminated AD and HC groups. Compartmental modeling confirmed that [^11^C]RO-643 had lower brain entry than both [^18^F]RO-963 and [^18^F]RO-948, and [^18^F]RO-948 showed a better contrast between (predicted) areas of high vs low tau accumulation. Thus, our subsequent analysis focused on [^18^F]RO-948. Both voxelwise and region-based analysis of [^18^F]RO-948 binding in HC vs AD revealed multiple areas where AD and HC significantly differed. Of 22 high-binding regions, 13 showed significant group difference (following ANOVA, F=45, p<10^-5^). Voxelwise analysis also revealed a set of symmetrical clusters where AD>HC (threshold of p<0.001, cluster size k>50).

**Conclusions:** [^18^F]RO-948 demonstrates superior characteristics to [^11^C]RO-643 and [^18^F]RO-963 for characterization of tau pathology in AD. Regional binding data and kinetic properties of RO-948 compare favorably with existing other tau PET tracers.

## INTRODUCTION

Building upon the success of in vivo imaging of amyloid protein aggregates in Alzheimer’s and related neurodegenerative disorders over the last decade, imaging of tau protein has become of scientific and medical interest. Alzheimer’s disease (AD), the most common of the “tauopathies” (*3*), is characterized by the presence of amyloid plaques and tau tangles at autopsy (*4,5*). The ability to image tau pathology in vivo is important for both differential diagnosis and in monitoring treatment response as new therapies include anti-tau mechanisms (*6–8*).

Recently, we reported the discovery and radiolabelling of several new candidate ligands for tau PET imaging (*1*). Through in vitro screening, autoradiography, and preclinical in vivo PET imaging studies (*2*), we selected three candidates for evaluation in AD dementia patients, as well as, younger and older control subjects (YC and OC, respectively) (*9*). The specificity of all three candidate tracers for tau aggregates was supported by autoradiography studies on AD Braak V/VI brain sections, co-localization of radiotracer binding with antibody staining of tau aggregates on the same tissue sections, and lack of co-localization with amyloid plaques. All three tracer candidates, ([^11^C]RO-963, [^11^C]RO-643 and [^18^F]RO-948 previously referred as [^11^C]RO6924963, [^11^C]RO6931643, and [^18^F]RO6958948, respectively), displayed good brain penetration as indicated by their peak standardized uptake value (SUV) and K_1_, rapid washout, and favorable metabolism in the non-human primate (*2*). The clinical evaluation of these three tracers and the selection of one, [^18^F]RO-948, for further evaluation is described below.

## MATERIALS AND METHODS

### Subjects

All subjects gave written informed consent before participation. The study protocol was reviewed and approved by the Johns Hopkins Medicine Institutional Review Board and registered on ClinicalTrials.gov (reference NCT0217627).

Three groups of participants were enrolled into the study: (1) YC, ages 25 – 40 years; (2) OC, aged over 50 years (healthy volunteers were recruited by PAREXEL International, Baltimore, MD); and (3) AD dementia subjects aged 50 years or older, recruited by Johns Hopkins. Screening consisted of routine medical assessments. AD dementia subjects met National Institute on Aging-Alzheimer’s Association (NIA/AA) diagnostic criteria, with Mini Mental State Exam (MMSE) scores of 16 – 26, inclusive. A positive amyloid PET scan was a prerequisite for AD subjects. Determination of capacity for consent was fulfilled (*10*). All subjects had a Body Mass Index (BMI) between 18-32, and if on dementia medication the dose was stable for at least four weeks prior to PET scanning.

### Study Design

The study aim was to directly compare the imaging properties of three novel tau PET tracers in human subjects and based on that comparison, select one tracer most suitable for application in clinical research for further evaluation.

The study was divided into three parts (Table 1):

**Table 1.** Study design and subject demographic.

| Part | Radioligands | <u>HC</u> |  |  | <u>AD</u> |  |  |
| --- | --- | --- | --- | --- | --- | --- | --- |
| | | Subjects | Plasma data (+) | Age ± SD | Subjects | Plasma data (+) | Age ± SD |
| **1** | [<sup>11</sup>C]RO-963<br>[<sup>11</sup>C]RO-643 | 2 (1F) | 2<br>2 | 30 ± 31 | 2 (1F) | 1<br>2 | 67 ± 69 |
| | [<sup>11</sup>C]RO-643<br>[<sup>18</sup>F]RO-948 | 2 (0F) | 2<br>2 | 25 ± 26 | 5 (2F) | 2<br>4 | 75.0 ± 7.8 |
| | [<sup>11</sup>C]RO-963<br>[<sup>18</sup>F]RO-948 | 3 (1F) | 3<br>2 | 32 ± 38 | 0 | | |
| **2A** | [<sup>18</sup>F]RO-948<br>Test – Retest | 5 (0F) | 4<br>4 | 62 ± 8 | 5 (1F) | 3<br>3 | 64.0 ± 6 |
| **2B** | [<sup>18</sup>F]RO-948<br>Dosimetry | 6 (3F) | NA | 58 ± 9 | | | |

Part 1 evaluated the kinetics of each of the three tracers in AD and YC. To avoid a possible confounding effect of inter-subject variability in tau burden (in AD subjects), a pairwise comparison was used in which each subject received two of the three tracers, with a washout of 7-14 days between the two scans (Table 1).

Subsequently, one tracer was selected for further evaluation in part 2A (test-retest imaging) and Part 2B (whole-body radiation dosimetry) (Table 1). Finally, four of the AD subjects who participated in Parts 1 (n=3) and 2A (n=1) returned for repeat [^18^F]RO-948 scans with no intervening tau or amyloid targeted treatment to provide a preliminary evaluation of the progression of tau deposition. The range of times between scans was 6.2 months to 21.6 months (mean 16.9 months).

### Radiochemistry

[^11^C]RO-963, [^11^C]RO-643, and [^18^F]RO-948 were prepared as described previously (*1*).

### Brain Imaging

A structural T1-weighted magnetic resonance brain scan was acquired on a Siemens Tim Trio 3T for all subjects other than dosimetry subjects. Data were acquired in the sagittal plane, using a 3D magnetization prepared rapid gradient echo (MP-RAGE) sequence with the following parameters: repetition time = 2.11 s, echo time = 2.73 ms, and inversion time = 1.1 s, and reconstructed in isotropic voxels of 0.8 mm. All MRI scans were inspected by a faculty neuroradiologist to exclude subjects with clinically relevant brain abnormalities. All brain PET imaging procedures were carried out at the Johns Hopkins Hospital PET center on a High-Resolution Research Tomograph (HRRT) (Siemens) with resolution of approximately 2 mm FWHM.

For brain PET scans, subjects were positioned in the tomograph after insertion of a venous cannula in an antecubital vein and a radial arterial cannula. The attenuation maps were generated from 6-minute transmission scans performed with a ^137^Cs point source before each radiotracer injection. Dynamic emission data were collected continuously for 0-90 minutes in Part 1 (frame durations: four 15-s, four 30-s, three 1-min, two 2-min, five 4-min, and twelve 5-min frames or a total of 30 frames for the 90-min scan. 5-min frames thereafter, for scans longer than 90 min) and up to 120 and 200 minutes in Parts 2A and 2B respectively, after intravenous bolus injection of [^11^C]RO-963, [^11^C]RO-643, or [^18^F]RO-948 in Part 1 and [^18^F]RO-948 in Part 2.

For radiation dosimetry (Part 2B), non-IV-contrast CT and PET images were obtained in four repetitions from the vertex to mid thighs. Images were acquired on a GE DVCT LySO-crystal 64-slice scanner. Each repetition of the vertex to mid thighs involved 8 bed positions over 91 minutes.

Target injected activities were as follows: 70-minute amyloid scans of 370 MBq (10 mCi) for ^18^F-Florbetapir (Amyvid) or 560 MBq (15 mCi) for [^11^C]-PIB dynamic PET scans; for 90 minutes scans of [^11^C]RO-963 and [^11^C]RO-643, 740 MBq (20 mCi); for 90-200 minute scans of [^18^F]RO-948, 185 MBq (5 mCi), later increased to up to 370 MBq (10 mCi); and for dosimetry scans of [^18^F]RO-948, 185 MBq (5 mCi).

Amyloid positivity was determined by FDA approved visual interpretation with [^18^F]-Florbetapir, or visual interpretation and SUV threshold with [^11^C]-PIB (*11*).

### Input function measurement

For all tau PET brain scans radial-arterial blood samples (optional for AD subjects) were obtained for derivation of a plasma input function (~42 samples over the duration of each tau PET brain scan), and for determination of radioactive metabolites in plasma with HPLC (6 or more samples). During the PET scan period, safety data were obtained including vital signs, ECG, labs, and safety monitoring.

### Outcome Measurements

Image data were reconstructed using the iterative ordered subsets expectation maximization algorithm, with correction for radioactive decay, dead time, attenuation, scatter and randoms (*12*).

Detailed analysis methods, including PET outcome variables are described in a companion paper *(13)*. Briefly, volumes of interest (VOIs) were defined by Freesurfer and FSL/FIRST software, manually refined based on each individual subjects’ MRI, and transferred to each subjects’ native PET space using the coregistration module of SPM12 (*14,15*) to generate time-activity curves (TACs) of VOIs. A total of 80 VOIs (left and right VOIs of 40 brain regions) were used. The VOI for the cerebellar cortex (Cb), served as a reference region, excluding the vermis and upper 6 mm to reduce contamination from the supra-tentorial regions. The main outcome variables were the blood-brain clearance rate constant (K_1_) from two-tissue compartmental analysis (TTCM), and the total distribution volume (V_T_) from plasma input graphical analysis (PRGA (*16*)) for Part 1; the distribution volume ratio (DVR) from PRGA, and the reference tissue graphical analysis (RTGA (*17*)), and standard uptake ratio (SUVR) to Cb for Part 2A. Part 2B outcome measures were activity time integrals for the whole-body dosimetry.

### [^18^F] RO-948 in AD and OC Subjects

The most promising radiotracer, [^18^F]RO-948, was further evaluated to compare regional tau accumulation in AD compared to OC. First, mean SUVR values of AD, OC, and YC subjects in all 80 regions were examined to explore the regional specific binding throughout the gray matter by ranking regions in order of mean SUVR values in AD subjects.

Next, regional SUVR data for the AD and OC groups were compared to examine the discriminatory power of the radiotracer.

Third, a voxel-wise analysis of differences in SUVR between AD and OC groups was performed with SPM12 (*14*). SUVR images of individual subjects were spatially normalized to the MNI space using parameters from each subjects’ PET-to-MRI co-registration and MRI-to-template spatial normalization given by respective modules of SPM12 (*15*) and smoothed by a Gaussian kernel of 6 mm full-width at half maximum (FWHM). A locally modified Desikan/Killiany atlas (*18*) of the 80 brain regions was used to identify the brain structures in MNI space. Separately, a SUVR threshold map (the mean SUVR map plus three times the median SD of gray matter voxels of OC subjects) was constructed for generation of a map of tau-positivity frequencies.

Subsequently, Braak stages (*19*) of tau accumulation were estimated using the region scheme and criteria of Schwarz et al. (*20*) which adapted the scheme of Braak (*19*) to SUVR images in the MNI space.

### Statistical analysis

To compare regional values of AD against OC we tested for the effect of region and group status (AD and OC) on PET outcome variables using analysis of variance (ANOVA) with group and region as the independent variables. If the ANOVA showed a significant effect of group, we used post-hoc t-tests, applying Bonferroni correction for multiple comparisons. Specifically, the Mann-Whitney U test was used because the outcome variables showed a skewed distribution. Statistical analyses were performed by Matlab.

In the exploratory voxel-wise statistical analysis, a significance threshold combination at p<0.001, uncorrected and k>50 (0.4 mL) was employed. The threshold for ‘commonly affected’ voxels in the tau-positive frequency map was set at the frequency value that roughly equated the resulting total volume of ‘commonly affected’ voxels to the total cluster volume of the SPM group difference analysis.

#### Whole body PET/CT Scanning

Mean residence times for each organ were calculated using the AUC of the non-decay corrected TAC and the tabulated organ volume as used in OLINDA/EXM® version 1.0 to calculate the radiation dose to individual organs, and effective dose equivalents and effective doses (ED).

## RESULTS

Subject demographics and study design are given in Table 1 and injected activity, mass and specific activity for all radiotracers (Supplementary Table S3). There were no serious adverse events.

### Brain uptake and kinetics

Peak brain SUV values were approximately 3.0, 1.5, and 3.5 for [^11^C]RO-963 (N=5), [^11^C]RO-643 (N=4), and [^18^F]RO-948, (N=5) respectively, for YC in the temporal lobe. Other regions showed a similar trend, with [^18^F]RO-948 having the highest peak SUV value, followed by [^11^C]RO-963, and then [^11^C]RO-643 (Figure 1). Brain penetration was highest for [^18^F]RO-948 and lowest for [^11^C]RO-643. The retention of [^11^C]RO-963 in YC at later times (60-90 min post injection) was somewhat higher than that of [^11^C]RO-643 and [^18^F]RO-948 suggesting that [^11^C]RO-963 has non-specific binding leading to its slower washout (Figure 1).

**Figure 1.**
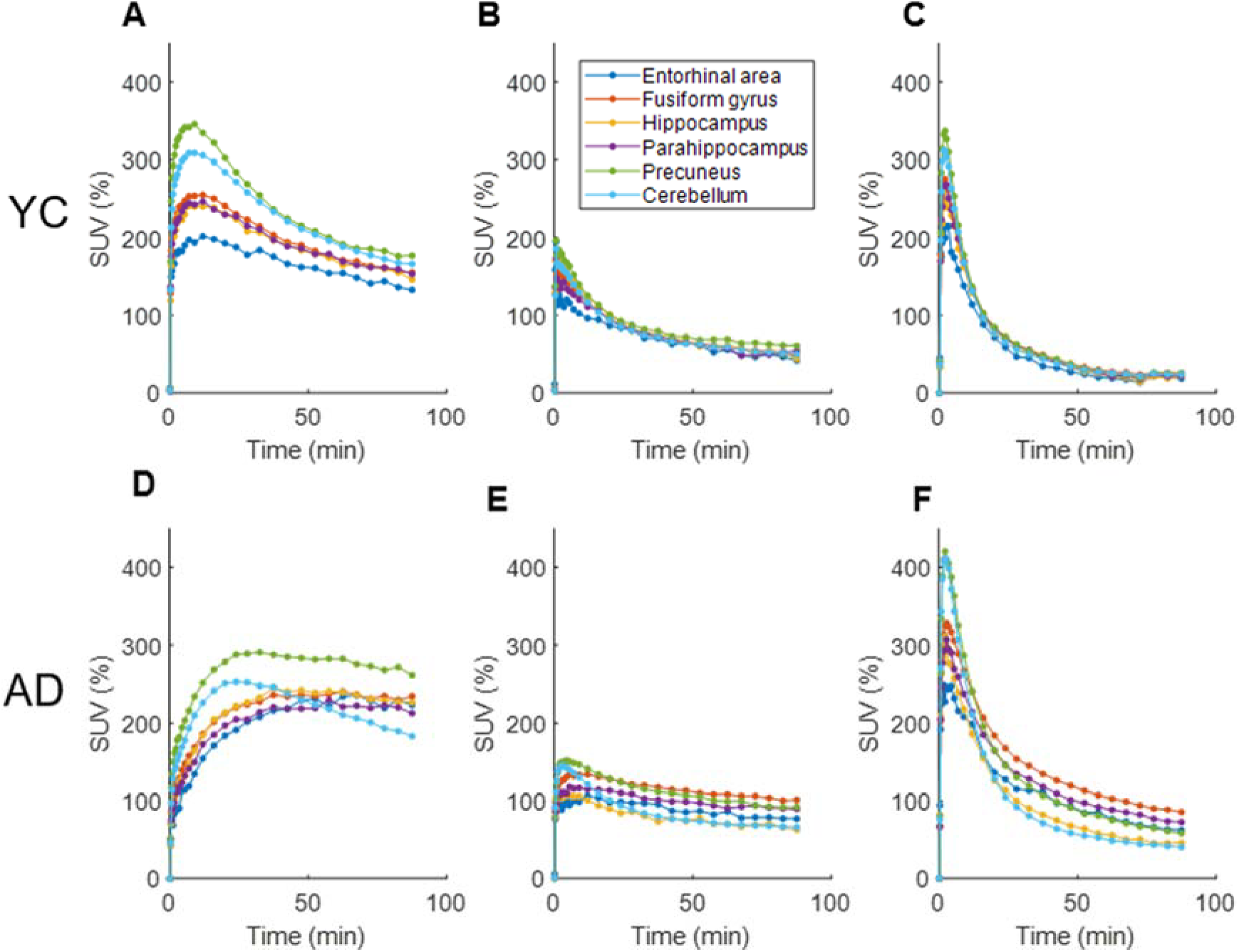
Line plots of time-activity curves (TACs) in standard uptake value (SUV) of selected brain regions of [^11^C]RO-963 (A,D), [^11^C]RO-643 (B,E), and [^18^F]RO-948 (C,F) for YC subjects (panels A-C) and AD, (panels D-F).

After direct comparison of brain uptake in the AD subjects of all 3 tracers with compartmental modeling (below), further testing of [^11^C]RO-963 was deprioritized.

When comparing [^11^C]RO-643 and [^18^F]RO-948 within AD subjects, [^18^F]RO-948 displayed a higher contrast between (predicted) tau-rich regions and tau-poor regions. The regional retention of [^18^F]RO-948 appeared to distinguish between subjects with AD and YC (Figure 2).

**Figure 2.**
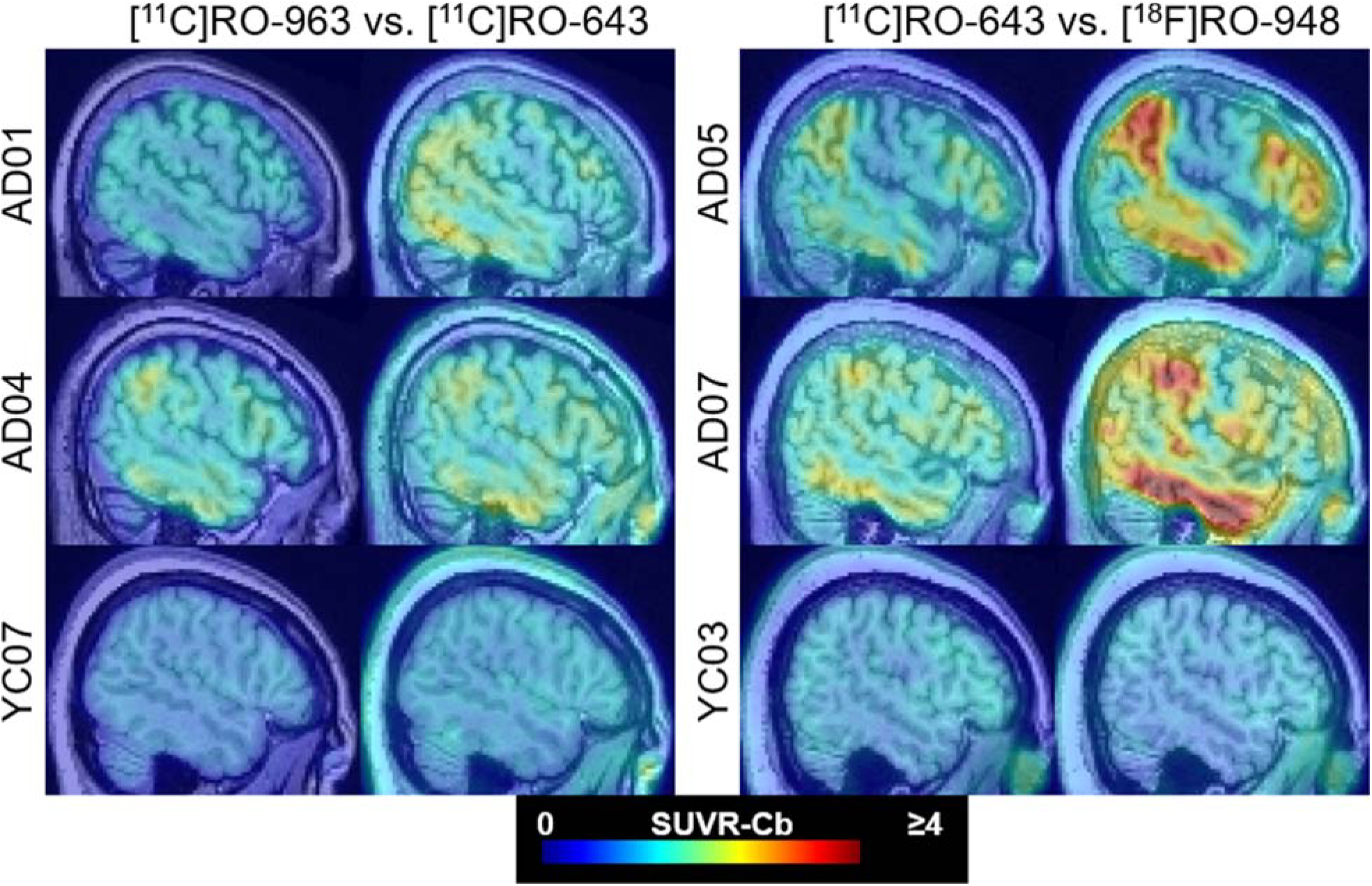
Sagittal SUVR images of the candidate radioligands, applied to the same AD and YC subjects.

TACs (60-90 min), for [^18^F]RO-948, for regions typically involved in tau accumulation (e.g., temporal lobe, fusiform, cingulate, parietal, occipital, parahippocampal) showed greater separation between the AD and YC groups than the other two tracers (Figure 1).

For comparison, the TACs for the cerebellar grey region are shown (Fig 1). Cerebellar grey matter is thought to contain low or no levels of aggregated tau in AD and YC subjects; thus, the late-phase tracer uptake should be low and similar between AD and YC subjects, best exhibited by [^18^F]RO-948.

Compartmental modelling (TTCM) confirmed the lower brain entry of [^11^C]RO-643 (mean AD) K_1_ = 0.054±0.01 and (YC) 0.065±0.02 mL/mL/min across regions) compared to [^11^C]RO-963 (mean (AD) K_1_ = 0.35±0.07 and (YC) 0.42±0.09 mL/mL/min) and [^18^F]RO-948 (mean (AD) K_1_ =0.35±0.1 and (YC) 0.43±0.9 and (OC) 0.42±0.14 mL/mL/min), a result anticipated from the TACs.

TACs for [^18^F]RO-948 YC compared to AD (Figure 1) are consistent with the anticipated low level of retention and regional heterogeneity in YC, and higher retention and regional heterogeneity in AD. The regional distribution of [^18^F]RO-948 retention was also distinct from that of the amyloid tracers (Figure 7).

**Figure 7.**
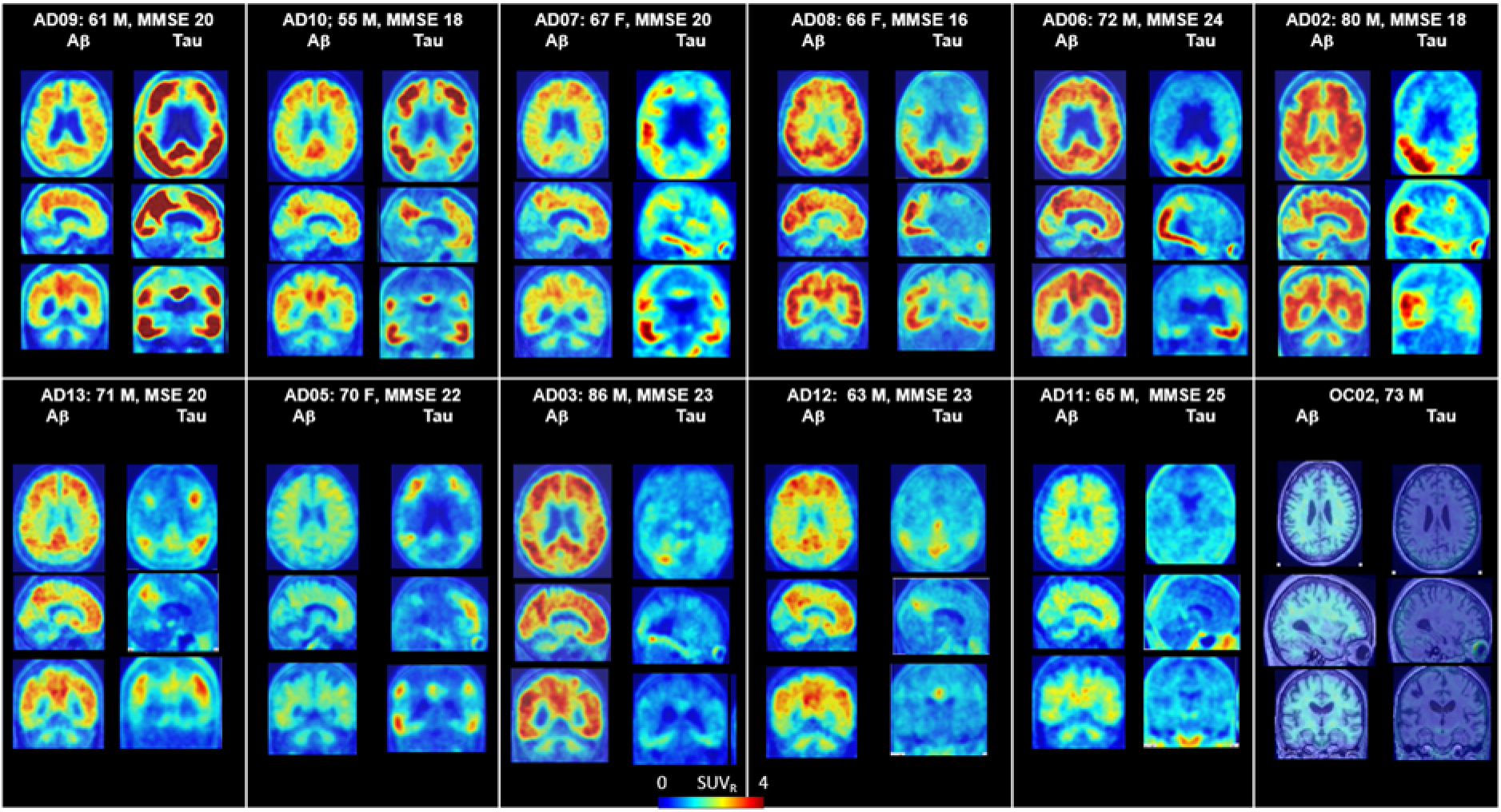
SUVR-Cb images of [^18^F]RO-948 of all 11 AD subjects, arranged in descending order of global mean SUVR, alongside the corresponding Aβ image ([^11^C]PiB SUVR, except for subjects AD02, AD05, and AD13 who had [^18^F]AV45 scans).

### Quantification

The kinetics of the three radiotracers was well described by a TTCM, and PRGA plots approached an asymptote by 20 min.

Regional (60-90 min) SUVR data (=y) correlated with DVR data for both [^11^C]RO-643 (y = 0.85·x + 0.22; R^2^ = 0.867; Figure 3A) and [^18^F]RO-948 (y = 1.62·x - 0.64; R^2^ = 0.911; 4 AD and 4 YC subjects *(13)*. Direct comparisons of SUVR data of [^11^C]RO-643 vs. [^18^F]RO-948 (Figure 3B) indicated a lower slope (0.36; 95% confidence intervals (CIs): 0.34 – 0.38) for AD data than YC data (slope = 0.51: 95% IC: 0.47 – 0.54). These data suggest that [^11^C]RO-643 underestimate SUVR in the AD range (i.e., the higher SUVR, the higher is the underestimation) compared to [^18^F]RO-948.

**Figure 3.**
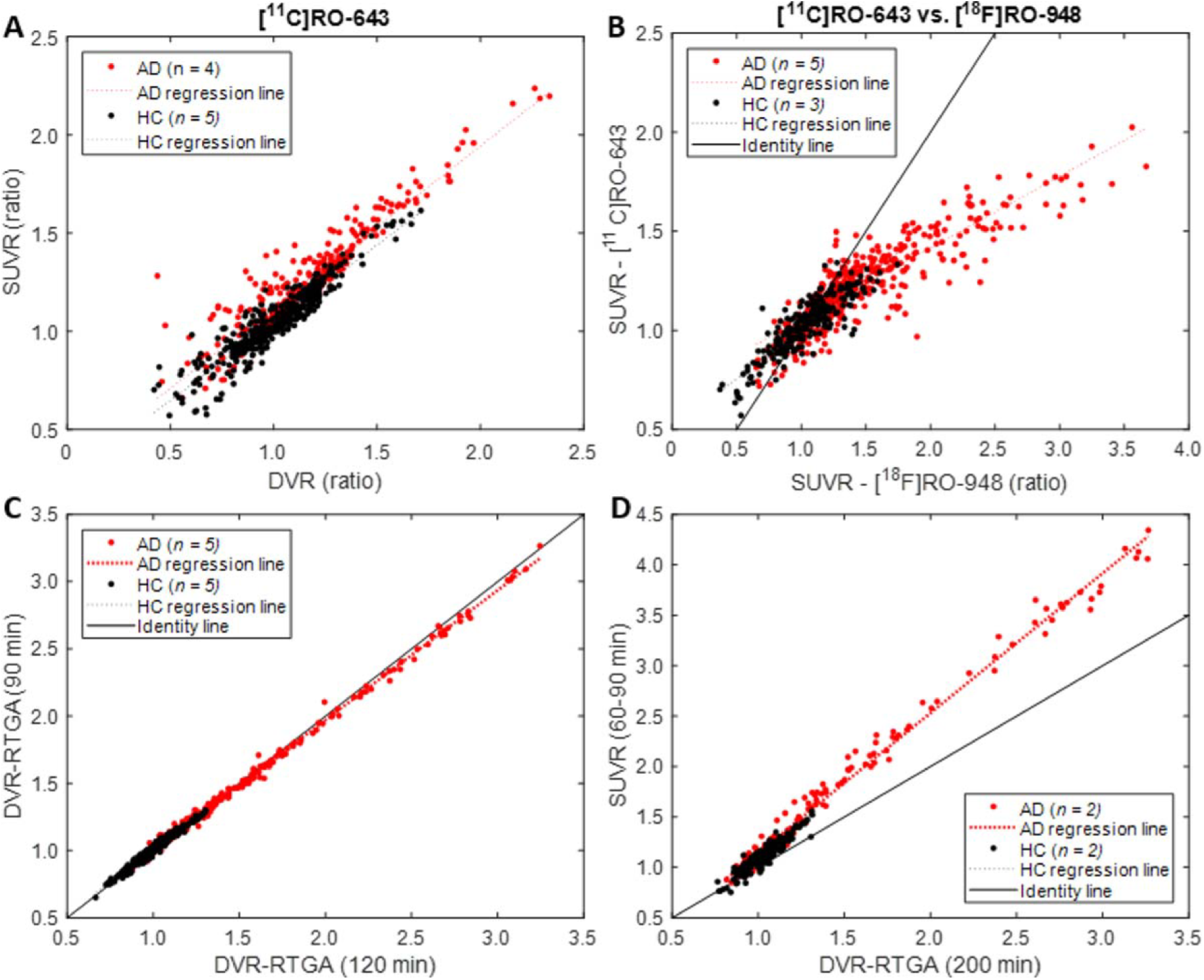
Scatter plots of SUVR (=y) versus DVR of [^11^C]RO-643 (A), and SUVR data of [^11^C]RO-643 (=y) versus [^18^F]RO-948 for subjects who had both scans (B). Scatter plots of DVR (RTGA), 90 versus 120 min circulation times for data analysis (C), and SUVR (=y) versus DVR (D) for [^18^F]RO-948.

Additionally, for [^18^F]RO-948 regional DVR data at 90 min (=y) linearly correlated with DVR data at 120 min for AD and OC subjects with a minimal bias of about 5% (Figure 3C) by reference tissue graphical analysis (RTGA; (*17*)). Note that RTGA yielded essentially identical DVR values to PRGA (considered the optimal method) for [^18^F]RO-948 (*13*). Regional SUVR values of 60–90 min frames were strictly proportional to DVR values of 200 min scans (Figure 3D, bottom right panel), which confirmed that SUVR of 60-90 min frames is a valid surrogate of DVR of 200 min scans for [^18^F]RO-948. Therefore, [^18^F]RO-948 was selected for further evaluation. A detailed evaluation of the kinetic modeling of [^18^F]RO-948 is given in the companion paper *(13)*.

### Injection Parameters

There were no significant differences in injected radioactivity, mass, or specific activity between test and retest studies of [^18^F]RO-948 (median range): Activity: test 271.6 MBq (166.5–364.8), retest 259.7 MBq (140.6–355.2), Student’s paired samples t-test (p=0.9); Injected mass:(test 0.08 µg (0.03–0.21), retest 0.13 µg (0.03–0.68), Student’s paired samples t-test (p=0.5); Specific activity: test 27.7 GBq/µmol (11–51), retest 33.0 GBq/µmol (16–71), Student’s paired samples t-test (p=0.5)).

### Metabolite analysis

[^11^C]RO-643 and [^18^F]RO-948 showed similar plasma parent time profiles (See Supplementary Figure S1). Across all subjects, the percentage of HPLC corrected radiolabeled [^18^F]RO-948 metabolites as compared to parent were very similar with no lipophilic metabolites identified. There was no evidence of defluorination of [^18^F]RO-948.

### Reproducibility of Blood Data Quantification

The percentage of the [^18^F]RO-948 parent compound in the plasma differed <10% between test and retest. For the first 5 minutes, test-retest differences was higher (25%), likely due to bolus injection differences (Supplementary Table S1).

### Reproducibility of PET Data Quantification

The test-retest variability with various methods (DVR, SUVR) in OC and AD test retest was 6-10% across regions *(13)*.

### Regional Uptake of [^18^F] RO-948

Mean SUVR values of the 40 regions (left and right merged) for AD, OC, and YC are shown in the supplemental material, Table S4 in descending order of mean SUVR of AD. Braak’s anterior regions (Hippocampus (Hp), Entorhinal Area (ER), and Parahippocampus (PH)) together with inferior temporal lobe (iTp), lateral cortical regions (inferior parietal lobe (iPa), supramarginal gyrus(SM), lateral occipital lobe (lOc), superior parietal lobe (sPa)), medial posterior regions (precuneus (Pr), posterior cingulate gyrus (pCg), isthmus/cingulate (IC)), and frontal regions (middle Frontal Lobe (mFr), rostral Frontal Lobe (rFr), frontal operculum (FO)) showed the highest SUVR values. Surface projection maps (Figure 4A) visually confirmed above-mentioned regional distributions of SUVR in AD.

**Figure 4.**
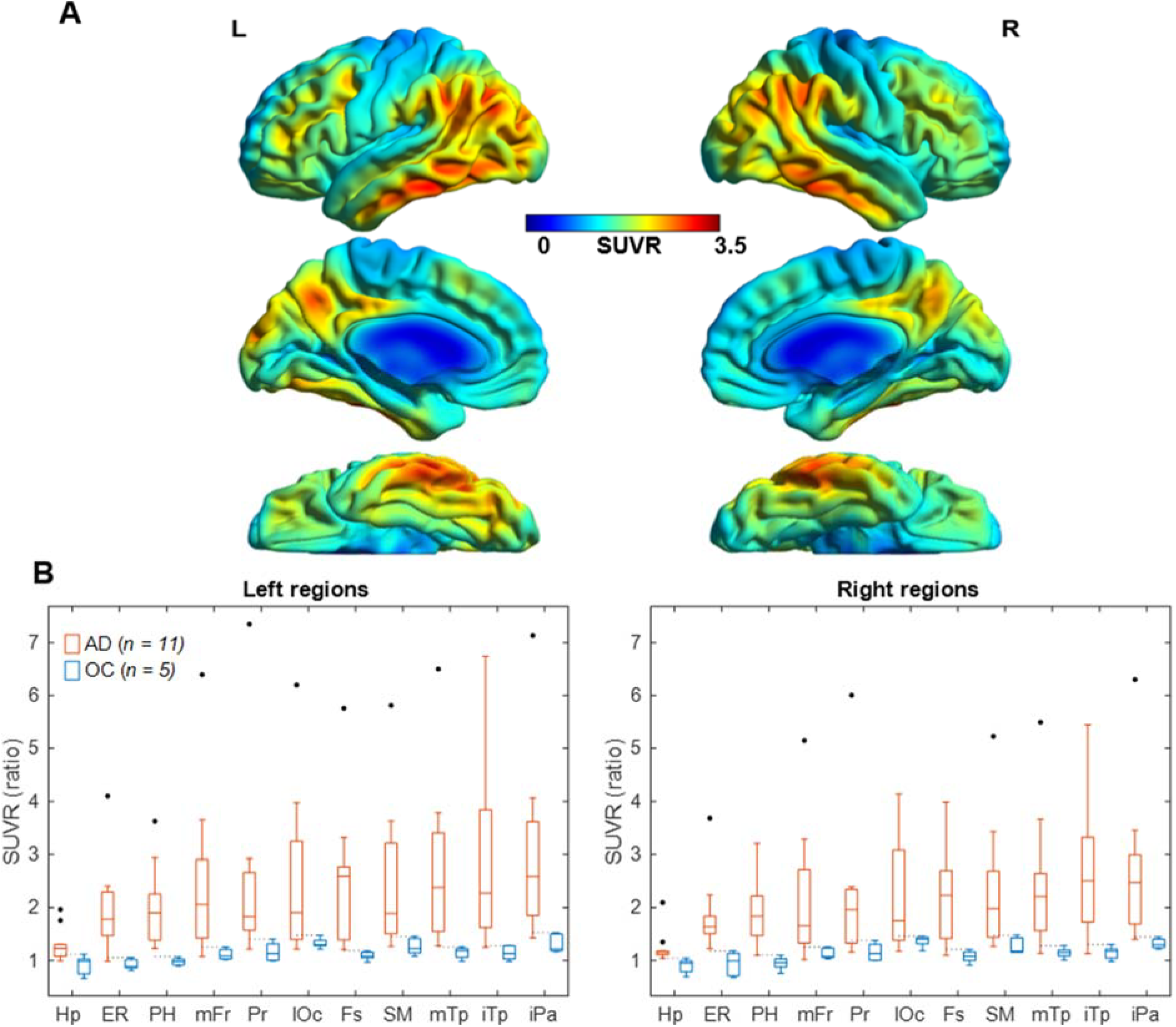
**A.** Surface projection maps of mean [^18^F]RO-948 SUVR images of AD subjects, lateral (upper), medial, and ventral (lower) views of the left (L) and right (R) hemispheres. **B.** Boxplot of SUVR data comparing AD and OC subjects in 8 regions in which AD subjects showed the highest mean SUVR values, and 3 Braak’s anterior regions (Hp, ER, and PH), left and right. Dots represent outlier points (i.e., outside ±2.7 SD, assuming normal distributions). See supplementary table S4 for full region names.

SUVR values of 22 regions, consisting of those showing high SUVR regions plus Braak’s anterior regions were compared between AD and OC in Figure 4B. The lowest AD values exceeded the highest OC values in 5 regions (3 left-sided and 2 right-sided; Table 2). Statistically, ANOVA showed a significant group effect (F=45, p<10^-5^), and 13 regions had significant differences (AD > OC) which survived Bonferroni correction (p uncorrected <0.05)/22=0.0023, Table 2).

**Table 2.** Presence (y) or absence (n) of separation (AD > maximal OC) and statistical differences (AD>OC; Mann-Whitney’s test).

|  |  | Hp | ER | PH | mFr | Pr | IOc | Fs | SM | mTp | iTp | iPa |
| --- | --- | --- | --- | --- | --- | --- | --- | --- | --- | --- | --- | --- |
| Separation | Left | n | n | y | n | n | n | y | n | y | n | n |
|  | Right | n | y | y | n | n | n | n | n | n | n | n |
| Statistical differences | Left | n | y | y | y | n | n | y | n | y | y | n |
|  | Right | y | y | y | n | n | n | y | n | y | y | y |

The voxel-wise statistical test (SPM12) identified a pair of roughly symmetrical clusters of AD>OC (Figure 5, A-D) that spanned Braak’s anterior regions and other high binding regions (iPa, and iTp) (Supplementary Materials Table S5). Along y-axis (anterior-posterior), both t-values (Supplementary Materials Table S5) and cluster volumes (Figure 5E) peaked around the anterior commissure (AC; y=0). If one selects a coronal slice of 1 cm thick that includes Hp (shown by green line in 5E) based on the SPM results, it is very likely such a slice coincides with Braak’s anterior block (blue horizontal lines) which was selected as a representative coronal slice to capture tau accumulations in AD (*19*).The map of Tau-positive frequencies had a similar volume of clusters when the threshold was set to 0.64 (i.e., 7 out of the 11 AD subjects). While extending to similar regions in the SPM analysis (Figure 5; panels F, G, and I correspond to panels A, B, and C, respectively), the clusters of the frequency map included Pc and pCg (panel J), and sPa and IPa (panels K, L).

**Figure 5.**
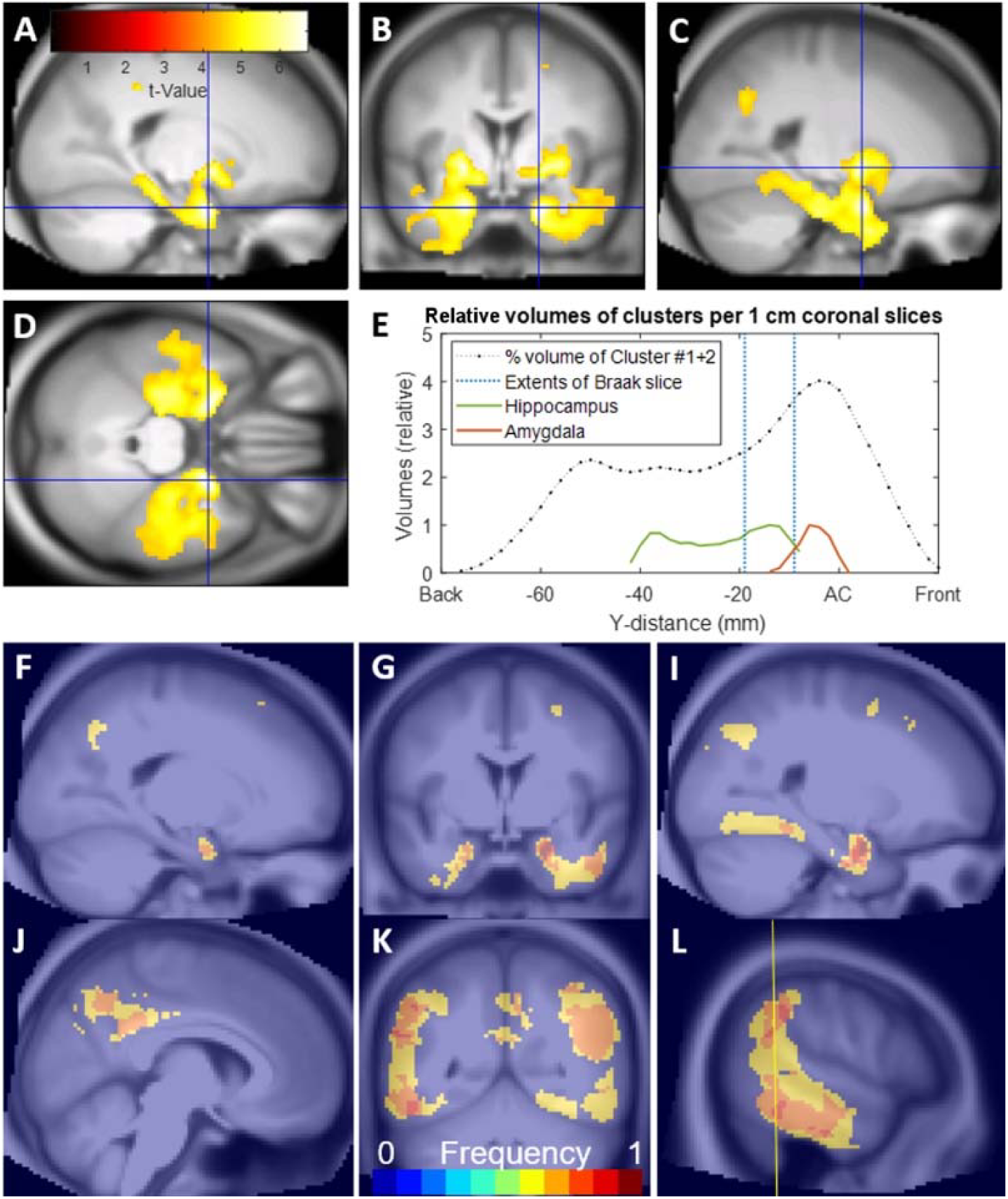
Results from SPM analysis (on std. MRI): A, C) Clusters #1 and 2, respectively. B,D) both lusters. E) Illustrates tau accumulation in AD in a representative coronal slice corresponding to Braak’s anterior block (19). Line plots of relative volumes of above-threshold voxels in 1 cm thick slices (the thickness of blocks of (19) and volumes of interest of the hippocampus and amygdala in 2 mm slices along the y-axis. Two vertical lines indicate the extents of the anterior slice of (19). Clusters of high [^18^F]RO-948 tau-positive frequencies (>7 of 11 AD subjects). F, G, I) SPM analysis of A, B and C. J, K, L) high agreement clusters in Pr, pCg, and IC (J), and iTp, SM, and iPa (K and L).

Tau-positivity results of Braak’s anterior and posterior regions are shown in Figure 6. Estimated Braak stages were Stage IV in 3 AD subjects (MMSE: 20, 22, 23), Stage V in 2 subjects (MMSE: 18, 20), and Stage VI in 6 subjects (MMSE: 16, 18, 20, 23, 24, 25). Single (Hp in 4 subjects) or multiple (3 subjects with -v) spared regions were noted across stages. No correlations of estimated Braak stages to MMSE scores were noted in this small samples of AD subjects (R^2^<0.01; p=0.86).

**Figure 6.**
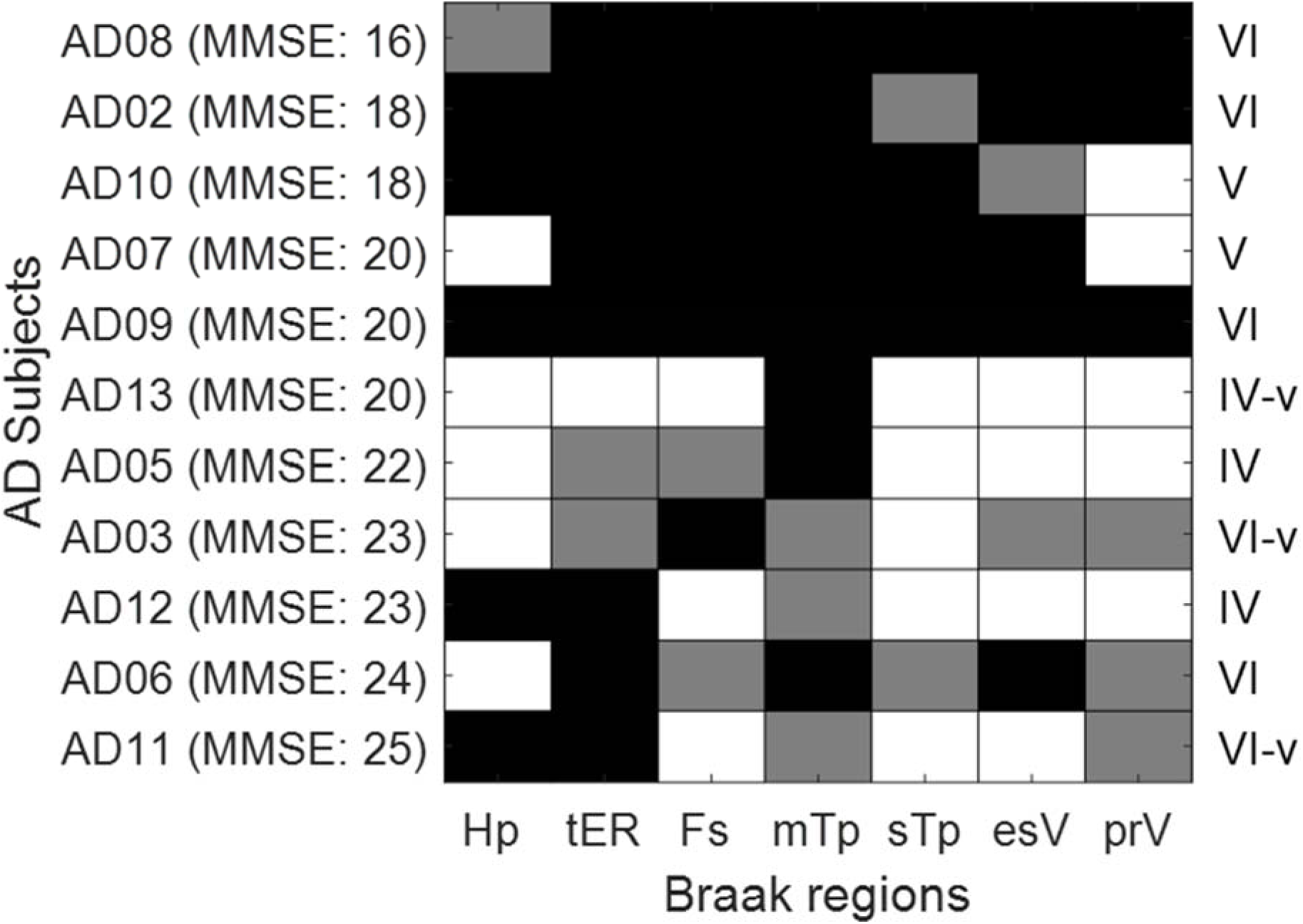
Checkerboard plot showing [^18^F]RO-948-positivity results (black cells: bilaterally positive; gray cells: unilaterally positive) of anterior and posterior Braak regions (Braak et al., 2006), and estimated Braak stages (right column) (Schwartz et al., 2016). Regions: hippocampus (HP), trans-entorhinal cortex (tER), fusiform gyrus (FS), and inferior, middle, and superior temporal cortices (iTp, mTp, and sTp), and the primary (prV) and extrastriatal (esV) visual cortices.

We illustrate amyloid load (using either [^11^C]PIB or [^18^F]AV45), Tau binding with [^18^F]RO-948, age and MMSE score in Figure 7 for all AD subjects. These figures show the ability of [^18^F]RO-948 to successfully image a range of Tau burden across a variety of patients and show the difference in regional patterns between amyloid and tau.

Finally, longitudinal studies in 4 AD subjects showed DVR/SUVR changes in 3 out of 4 cases over one year with considerable increases in Tau in regions selected a priori (Supplemental figure S2). These data suggest that [^18^F]RO-948 could be used to detect changes of Tau pathology over time.

### Whole Body Radiation Dosimetry

All organs TACs were well fit with 1-2 exponential functions. The gallbladder received the highest dose of 0.15 mSv/MBq (0.5 rem/mCi) with most organs receiving 0.01-0.05 mSv/MBq (0.03–0.15 rem/mCi).

The mean effective dose (averaged for males and females) was estimated to be 0.015 mSv/MBq (0.056 rem/mCi). Detailed organ dosimetry is given in supplemental material (Table S2).

## DISCUSSION

We report the in vivo characterization of 3 novel tau tracers [^11^C]RO-963, [^11^C]RO-643, and [^18^F]RO-948 using an arterial input. Due to its superior in vivo imaging properties, [^18^F]RO-948 was selected for further evaluation. Healthy subjects demonstrated minimal retention, while AD dementia subjects demonstrated regional distributions consistent with published post-mortem data on PHF-tau.

Kinetic analyses demonstrated good correlations and time stability for measures by the two-tissue compartmental model and PRGA. DVR (PRGA) remained constant between 180 and 200 min (*see companion paper Kuwabara et al.*). DVR (RTGA) correlated with DVR of PRGA, suggesting that RTGA using the cerebellum grey as the reference tissue could be utilized for dynamic studies. SUVR-Cb and DVR showed excellent test-retest variability results in the small sample of patients evaluated to date. SUVR-Cb (60-90 min) correlates linearly with DVR. Altogether, above findings demonstrated SUVR-Cb was suitable for larger clinical studies. In fact, SUVR-Cb demonstrated excellent separation between AD and OC in twelve selected regions.

The human whole-body dosimetry is lower or comparable to that of other [^18^F] tau radiotracers and allows as many as 6 scans per subject at 10 mCi (370 MBq) each using US guidelines.

In common with flortaucipir ([^18^F]AV1451, T807), we observed the highest uptake of [^18^F]RO-948 in inferior and lateral temporal lobes of AD patients (*21*), and in other brain regions known to display tau pathology in AD. In contrast to flortaucipir, [^18^F]RO-948 shows stable SUVRs in AD subjects at around 70-80 min. This may confer some advantages to the use of [^18^F]RO-948 in longitudinal studies or those assessing the impact of a therapeutic agent. In this limited dataset we saw no appreciable uptake in the choroid plexus and in the striatum. Off-target retention was observed in the substantia nigra and, in one control subject, in the cerebellar vermis. Extracerebral tracer retention was occasionally observed in the meninges and more often in the retina.

Recently a number of other tau tracers have been reported including: [^18^F]THK5351 (*22*), [^18^F]MK6240 (*23,24*), [^18^F]GTP1 (*25*), [^18^F]PI2620 (*26*), and [^18^F]JNJ643493 (*27*). When comparing these tracers to each other, and [^18^F]RO-948, assuming no major differences in selectivity (e.g. a large fraction of the specific signal of [^18^F]THK5351 has been shown to be attributable to MAO-B rather than tau (*28*)), or kinetics, a number of other factors should be considered, particularly if the intended application of the tracer is in longitudinal and or multicenter studies. First, production needs to be reliable and consistently have a sufficiently high yield (as in the case of [^18^F]RO-948) for it to be feasible for the tracer to be transported between production and scanning sites. Second, extra-cerebral uptake should be minimal to avoid spill over into the cortex complicating quantification and interpretation of images, particularly in subjects with mild AD. Finally, a high specific signal may increase tracer sensitivity to detect low levels of pathology in earlier stages of the disease and to appreciate pathology changes over time. [^18^F]RO-948 satisfies these criteria. The tau-positive frequency map, advanced by Cho et al (*29*), is an example of qualitative assessment (spatial occurrences of above-threshold voxels). Such maps are expected to supplement voxel wise statistical tests or quantitative assessments. With greater numbers of subjects (67 OC and 53 AD subjects), the paper reported a maximal frequency of 0.3, while this study with [^18^F]RO-948 showed much higher frequencies (≥0.7) in similar regions. The difference could be ascribed to frequencies of subtypes of AD in the two studies. In fact, frequency maps could supplement mean SUVR maps of subtypes of AD (*30*) the latter of which could be influenced by subjects with high SUVR values. Clearly a much larger sample of [^18^F]RO-948 OC and AD subjects are needed to confirm this observation of frequency differences.

Currently Braak’s stages are applied to tau-neuroimaging studies either using a conventional approach involving the whole brain regions (*20,31,32*) or using a new approach using anterior and posterior brain blocks alone (*19,20*). By either approach, studied AD subjects tended to be in either stage V or VI (62% by Cho; 87% by Scholl; 78% by Schwarz; 71% by this study). However, it should be noted that there are several reasons why one would not expect tau PET images to correspond directly to the Braak system. Perhaps most importantly, Braak staging is based on counting of NFT lesions identified on post-mortem tissue. In contrast a PET tracer binding to tau fibrils will presumably reflect the density of a wider range of tau aggregates. Therefore, new staging algorithms are needed to capture natural advancement of tau-accumulation, differences in tau-accumulation patterns in subtypes of AD, and treatment effects of tau-targeting treatments. Sparing of selected regions, as proposed by Schwarz et al. (*20*) may be one possibility, as we also observed sparing of regions with [^18^F]RO-948 in this study. These findings are consistent with the heterogeneity of binding for all of the recent tau radiotracers and the need for novel classifications.

Our limited sample of longitudinal data suggests that [^18^F]RO-948 is able to detect longitudinal Tau changes (*33*) though larger longitudinal studies are needed to confirm its reliability and sensitivity to track tau pathology changes over time.

## CONCLUSIONS

[^18^F]RO-948 is a promising radiotracer imaging tau pathology in AD. It shows good brain uptake, has no apparent brain penetrant radio-labelled metabolites, has a good kinetic profile, shows little or no retention in cognitively normal young or elderly control subjects and a distribution in AD dementia subjects consistent with published post-mortem data, has low test-retest variability, and radiation dosimetry consistent with that of other ^18^F-labelled CNS PET tracers.

It is our hope that tools such as [^18^F]RO-948 will allow us to gain a better understanding of the pathophysiology of AD, and in the context of drug development select patients for clinical trials, confirm the mechanism of action of drugs targeting pathological tau, and monitor the effects of disease modifying therapies regardless of whether or not they target tau directly.

## Acknowledgements

Special thanks to the Johns Hopkins PET center staff and dosimetry calculations by Michael Stabin, Vanderbilt University.

The study was funded by F. Hoffmann-La Roche, Basel, Switzerland.

SMR and MT are supported by the Intramural Research Program, National Institute on Aging, NIH.

## Reference list

1. Gobbi LC, Knust H, Korner M, et al. Identification of Three Novel Radiotracers for Imaging Aggregated Tau in Alzheimer’s Disease with Positron Emission Tomography. Journal of Medicinal Chemistry. 2017;60(17):7350–7370

2. Honer M, Gobbi L, Knust H, et al. Preclinical Evaluation of [18]F-RO6958948, [11]CRO6931643 and [11 C]-RO6924963 as Novel Radiotracers for Imaging Aggregated Tau in Alzheimer’s Disease with Positron Emission Tomography. Journal of Nuclear Medicine. 2017 Sep 28. [Epub ahead of print]

3. Villemagne VL, Okamura N. Tau imaging in the study of ageing, Alzheimer’s disease, and other neurodegenerative conditions. Current opinion in neurobiology. 2016;36:43–51.

4. Hyman BT, Phelps CH, Beach TG, et al. National Institute on Aging-Alzheimer’s Association guidelines for the neuropathologic assessment of Alzheimer’s disease. Alzheimer’s & dementia. 2012;8:1–13.

5. Murayama S, Saito Y. Neuropathological diagnostic criteria for Alzheimer’s disease. Neuropathology. 2004;24:254–260.

6. Anand K, Sabbagh M. Early investigational drugs targeting tau protein for the treatment of Alzheimer’s disease. Expert opinion on investigational drugs. 2015;24:1355–1360.

7. Panza F, Solfrizzi V, Seripa D, et al. Tau-Centric Targets and Drugs in Clinical Development for the Treatment of Alzheimer’s Disease. BioMed Research International, 2016: Article ID 3245935, doi:10.1155/2016/3245935

8. Schroeder SK, Joly-Amado A, Gordon MN, Morgan D. Tau-Directed Immunotherapy: A Promising Strategy for Treating Alzheimer’s Disease and Other Tauopathies. Journal of NeuroImmune Pharmacology. 2016;11:9–25.

9. Wong DF, Borroni E, Kuwabara H, et al. First in-human PET study of 3 novel tau radiopharmaceuticals: [^11^C]RO6924963, [^11^C]RO6931643, and [^18^F]RO6958948. Alzheimer’s & Dementia.2015; 11:P850–P851.

10. Alzheimer’s Association. Research consent for cognitively impaired adults: recommendations for institutional review boards and investigators. Alzheimer disease and associated disorders. 2004;18:171–175.

11. Ng S, Villemagne VL, Berlangieri S, et al. Visual assessment versus quantitative assessment of [^11^C]-PIB PET and [^18^F]-FDG PET for detection of Alzheimer’s disease. Journal of nuclear medicine. 2007;48:547–552.

12. Rahmim A, Cheng J-C, Blinder S, Camborde M-L, Sossi V. Statistical dynamic image reconstruction in state-of-the-art high-resolution PET. Physics in Medicine and Biology. 2005;50:4887–4912.

13. Kuwabara H, Comley RA, Borroni E, et al. Evaluation of [18F]RO-948 ([18F]RO6958948) for quantitative assessment of tau accumulation in the human brain with positron emission tomography. 2017 *[submitted as companion paper to Journal of Nuclear Medicine, Volume, Issue, etc to be filled in if accepted]*

14. Friston KJ. (2003) Introduction: Experimental design and statistical parametric mapping. In Frackowiak RSJ, Friston KJ, Frith C, Dolan R, Price CJ, S. Zeki, Ashburner J, and Penny WD, editors, Human Brain Function. Academic Press, 2nd edition, pp 599–634

15. Ashburner J, Friston KJ. Rigid body registration. In: Frackowiak RSJ, Friston KJ, Frith C, et al., eds. Human Brain Function. 2nd ed: Academic Press; 2003.

16. Logan J, Fowler JS, Volkow ND, et al. Graphical analysis of reversible radioligand binding from time-activity measurements applied to [N-^11^C-methyl]-(-)-cocaine PET studies in human subjects. Journal of cerebral blood flow and metabolism 1990;10:740–747.

17. Logan J, Fowler JS, Volkow ND, Wang GJ, Ding YS, Alexoff DL. Distribution volume ratios without blood sampling from graphical analysis of PET data. Journal of cerebral blood flow and metabolism. 1996;16:834–840.

18. Desikan RS, Ségonne F, Fischl B, et al. An automated labeling system for subdividing the human cerebral cortex on MRI scans into gyral based regions of interest. Neuroimage. 2006;31:968–980.

19. Braak H, Alafuzoff I, Arzberger T, Kretzschmar H, Del Tredici K. Staging of Alzheimer disease-associated neurofibrillary pathology using paraffin sections and immunocytochemistry. Acta Neuropathologica. 2006;112:389–404.

20. Schwarz AJ, Yu P, Miller BB, et al. Regional profiles of the candidate tau PET ligand [^18^F]-AV-1451 recapitulate key features of Braak histopathological stages. Brain 2016;139:1539.

21. Pontecorvo MJ, Devous MD, Navitsky M, et al. Relationships between flortaucipir PET tau binding and amyloid burden, clinical diagnosis, age and cognition. Brain: A Journal of Neurology. 2017;140:748–763.

22. Harada R, Ishiki A, Kai H, et al. Correlations of [^18^F]-THK5351 PET with post-mortem burden of tau and astrogliosis in Alzheimer’s disease. Journal of nuclear medicine. 2017 Sept 1 [Epub ahead of print].

23. Hostetler ED, Walji AM, Zeng Z, et al. Preclinical Characterization of [^18^F]-MK-6240, a Promising PET Tracer for In Vivo Quantification of Human Neurofibrillary Tangles. Journal of nuclear medicine. 2016;57:1599–1606.

24. Walji AM, Hostetler ED, Selnick H, et al. Discovery of 6-(Fluoro-(18)F)-3-(1H-pyrrolo[2,3-c]pyridin-1-yl)isoquinolin-5-amine ([(18)F]-MK-6240): A Positron Emission Tomography (PET) Imaging Agent for Quantification of Neurofibrillary Tangles (NFTs). Journal of Medicinal Chemistry. 2016;59:4778–4789.

25. Sanabria Bohorquez S, Barret O, Tamagnan G, et al. Assessing optimal injected dose for tau PET imaging using [^18^F]GTP1 (Genentech Tau Probe 1). Journal of Nuclear Medicine. 2017;58:848.

26. Barret O, Seibyl J, Stephens A, et al. First-In-Human PET Studies With The Next Generation Tau Agent 18-F PI-2620 In Alzheimer’s Disease, Progressive Supranuclear Palsy, and Controls. Alzheimer’s & Dementia. 2017;13:P3–P4.

27. Declercq L, Rombouts F, Koole M, et al. Preclinical Evaluation of [^18^F]-JNJ64349311, a Novel PET Tracer for Tau Imaging. Journal of nuclear medicine. 2017;58:975–981.

28. Ng KP, Pascoal TA, Mathotaarachchi S, et al. Monoamine oxidase B inhibitor, selegiline, reduces (18)F-THK5351 uptake in the human brain. Alzheimer’s Research & Therapy. 2017;9:25.

29. Cho H, Choi JY, Hwang MS, et al. In vivo cortical spreading pattern of tau and amyloid in the Alzheimer disease spectrum. Annals of Neurology. 2016;80:247–258.

30. Ossenkoppele R, Schonhaut DR, Scholl M, et al. Tau PET patterns mirror clinical and neuroanatomical variability in Alzheimer’s disease. Brain. 2016;139:1551–1567.

31. Scholl M, Lockhart SN, Schonhaut DR, et al. PET Imaging of Tau Deposition in the Aging Human Brain. Neuron. 2016;89:971–982.

32. Braak H, Braak E. Neuropathological stageing of Alzheimer-related changes. Acta Neuropathoogica. 1991;82:239–259.

33. Kuwabara H, Borroni E, Comley RA, et al. On Evaluation Of Tau Accumulations In Longitudinal Studies Of Alzheimer’s Disease (AD): Implications From a PET Study With [18F]RO6958948. Alzheimer’s & Dementia. 13:P139.

